# Single-cell screening of SARS-CoV-2 target cells in pets, livestock, poultry and wildlife

**DOI:** 10.1101/2020.06.13.149690

**Authors:** Dongsheng Chen, Jian Sun, Jiacheng Zhu, Xiangning Ding, Tianming Lan, Linnan Zhu, Rong Xiang, Peiwen Ding, Haoyu Wang, Xiaoling Wang, Weiying Wu, Jiaying Qiu, Shiyou Wang, Haimeng Li, Fuyu An, Heng Bao, Le Zhang, Lei Han, Yixin Zhu, Xiran Wang, Feiyue Wang, Yuting Yuan, Wendi Wu, Chengcheng Sun, Haorong Lu, Jihong Wu, Xinghuai Sun, Shenghai Zhang, Sunil Kumar Sahu, Haixia Chen, Dongming Fang, Lihua Luo, Yuying Zeng, Yiquan Wu, ZeHua Cui, Qian He, Sanjie Jiang, Xiaoyan Ma, Weimin Feng, Yan Xu, Fang Li, Zhongmin Liu, Lei Chen, Fang Chen, Xin Jin, Wei Qiu, Huanming Yang, Jian Wang, Yan Hua, Yahong Liu, Huan Liu, Xun Xu

## Abstract

A few animals have been suspected to be intermediate hosts of severe acute respiratory syndrome coronavirus 2 (SARS-CoV-2). However, a large-scale single-cell screening of SARS-CoV-2 target cells on a wide variety of animals is missing. Here, we constructed the single-cell atlas for 11 representative species in pets, livestock, poultry, and wildlife. Notably, the proportion of SARS-CoV-2 target cells in cat was found considerably higher than other species we investigated and SARS-CoV-2 target cells were detected in multiple cell types of domestic pig, implying the necessity to carefully evaluate the risk of cats during the current COVID-19 pandemic and keep pigs under surveillance for the possibility of becoming intermediate hosts in future coronavirus outbreak. Furthermore, we screened the expression patterns of receptors for 144 viruses, resulting in a comprehensive atlas of virus target cells. Taken together, our work provides a novel and fundamental strategy to screen virus target cells and susceptible species, based on single-cell transcriptomes we generated for domesticated animals and wildlife, which could function as a valuable resource for controlling current pandemics and serve as an early warning system for coping with future infectious disease threats.

## Introduction

In the past two decades, the world has witnessed the outbreak and spread of SARS, Middle East respiratory syndrome (MERS)^1^, ZIKA^2^, avian influenza and swine influenza^3^, which have been posing an urgent challenge to our infectious disease prevention and control system. Recently, SARS-CoV-2 has caused a highly contagious pandemic disease named coronavirus disease 2019 (COVID-19), which is rapidly spreading all over the world and has triggered a severe public health emergency. As of 3rd June 2020, globally the total number of confirmed COVID-19 cases and deaths has uncontrollably reached 6,287,771 and 379,941, respectively^4^. The bat has been proposed to be the original host of SARS-CoV-2^5^, however, the transmission from bats to humans requires some intermediate hosts. Several studies have linked pangolins, cats, dogs and hamsters with SARS-CoV-2 infection and transmission^6–12^, indicating the potential widespread prevalence across animals, which would post potential threats to humans. The identification of the origin of this virus and its path to becoming a deadly human pathogen is needed to understand how such processes occur in nature and identify ways we can prevent the onset of these types of global crises in the future.

Evaluating host susceptibility is critical for controlling the infectious disease. The screening of virus putative host is usually performed using in vivo assay or inoculation experiments, which helps to reveal the host susceptibility, however, there are several limitations: 1) Experiments for some dangerous and infectious viruses need to be performed in a biosafety level 3 or level 4 laboratories, meaning limited number of researchers or groups can participate in the host screening work. 2) Only limited types of viruses and limited number of animals can be evaluated each time, thus the screening throughput is relatively low. Host range of a virus is closely associated with the availability of virus receptors, thus understanding the expression patterns of virus entry factors is of fundamental importance, and could play pivotal role in controlling the virus spread in current and future pandemics. Determining the target cells of SARS-CoV-2 based on the relative expression of virus entry factors provides potential clues to narrow down the putative intermediate hosts. The entry of SARS-CoV-2 into host cell is initiated by the binding of virus spike glycoprotein (S) to cell receptor angiotensin-converting enzyme 2 (*ACE2*)^13^ and the cleavage of S protein by transmembrane serine protease 2 (*TMPRSS2*)^14^. Although SARS-CoV-2 like corona virus has been isolated from pangolins and bats, their susceptible cell types for SARS-CoV-2 is not clear. Given that the species barrier of SARS-CoV-2 was estimated to be relatively low^13^ and livestock, poultry and pets have very close contact with humans, it is crucial to evaluate animal susceptibility to SARS-CoV-2. Previous studies have proposed that animal tissues show high heterogeneity in terms of cellular composition and gene expression profiles^15^, and *ACE2* is only expressed in a small proportion of specific cell populations^16^, making single cell analysis of SARS-CoV-2 target cells an attracting field to investigate. Here, we constructed the single cell atlas for livestock, poultry, pets and wildlife, then screened putative SARS-CoV-2 target cells (indicated by the co-expression patterns of SARS-CoV-2 entry receptor *ACE2* and SARS-CoV-2 entry activator *TMPRSS2*) and systematically evaluated their susceptibility, with the aim to understand the virus transmission routes and provide clues to fight against COVID-19.

## Results

### Construction of the single-cell atlas for different tissues of pangolin, cat and pig

Both pangolin and cat are suspected to be SRAS-CoV-2 intermediate hosts. SRAS-CoV-2 like coronavirus has been isolated from pangolin, and SARS-CoV-2 was proposed to origin from the recombination of a pangolin coronavirus with a bat coronavirus^6,7^. Cat is also a suspected intermediate host, as human-to-cat and cat-to-cat transmission of SARS-CoV-2 have been reported^9,10^. Domestic pig is an animal in close contact with human and have been reported to be susceptible to SARS coronavirus^17^. Although those animals have been linked with coronavirus, yet a comprehensive single-cell atlas for those species is missing. In this study, we generated the single nuclei libraries for various tissues of pangolin (heart, liver, spleen, lung, kidney, large intestine, duodenum, stomach and esophagus), cat (heart, liver, lung, kidney, eyelid, esophagus, duodenum, colon and rectum), and pig (heart, liver, spleen, lung, kidney, hypothalamus, area postrema, vascular organ of lamina terminalis, subfomical organ and cerebellum) (Fig. 1a, Supplementary Table 1). In total, 99740, 35345 and 92863 single cell transcriptomes passing quality control (see methods) were obtained for cat, pangolin and pig respectively (Fig. 1b-j, Supplementary Table 1). Cell clustering were performed using Seurat^18,19^ and cell type annotation were conducted according to cluster differentially expressed genes (DEGs) and the expression of canonical cell type markers (Extended Data Fig. 1–3, Supplementary Table 2). Overall, the high quality and comprehensive single cell atlas for distinct organs of three coronavirus susceptible animals were generated in this study, which provides valuable resources for further studies of their cellular taxonomy and makes it possible to identify virus target cells and screen host susceptibility at single cell level.

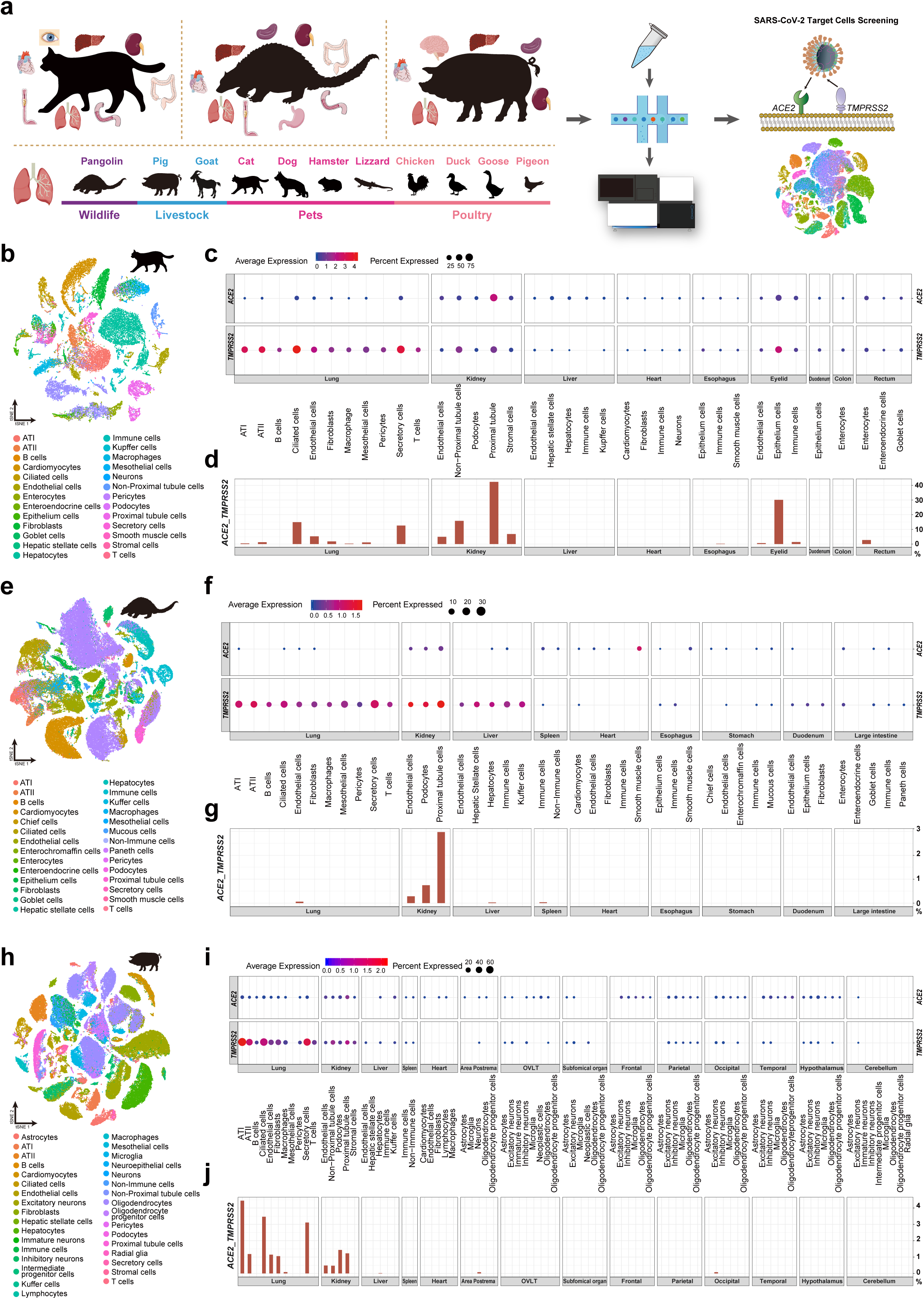

### Construction of the single-cell atlas for lungs of livestock, poultry, pets and wildlife

Lung is one of the main target organ of SARS-CoV-2^20^ and pneumonia is the typical symptom of COVID-19^21^. To evaluate the detailed expression patterns of *ACE2* and *TMPRSS2* in lung cells for various species, we generated single nuclei libraries for livestock (pig, goat), poultry (chicken, pigeon, goose, duck), pets (cat, dog, hamster, lizard) and wildlife (pangolin), resulting in a total of 123,445 cells passing quality control (Supplementary Table 1). In total, eleven cell types (ATI, ATII, ciliated cells, secretory cells, endothelial cells, fibroblasts, mesothelial cells, pericytes, T cells, B cells and macrophages) were identified in comparative lung atlas (Extended Data Fig. 4, Supplementary Table 2).

### Screening of SARS-CoV-2 target cells in different organs of cat, pangolin and pig

In cat, *ACE2* and *TMPRSS2* co-expressing cells were detected in lung (ATI, ATII, secretory cells, mesothelial cells, ciliated cells, endothelial cells, fibroblasts, macrophages), kidney (endothelial cells, non-proximal tubule cells, proximal tubule cells, stromal cells); eyelid (endothelial cells, epithelium cells and immune cells), esophagus (immune cells) and rectum (enterocytes). Notably, we observed over 40% co-expression of *ACE2* and *TMPRSS2* in proximal tubule cells of cat kidney, and around 30% in epithelium cells of cat eyelid (Fig. 1d, Supplementary Table 3).

In pangolin, SARS-CoV-2 target cells were found in lung endothelial cells, kidney (endothelial cells, podocytes and proximal tubule cells), liver (hepatocytes) and spleen (immune cells) (Fig. 1g, Supplementary Table 3).

In pig, *ACE2* and *TMPRSS2* were mainly co-expressed in lung (ATI, ATII, ciliated cells, secretory cells, endothelial cells, fibroblasts, macrophages) and kidney (non-proximal tubule cells, proximal tubule cells, endothelial cells, podocytes) (Fig. 1j, Supplementary Table 3). Overall, the proportions of SARS-CoV-2 target cells in cat are much higher than the proportions in corresponding cell types of all other species studied here.

### Screening of SARS-CoV-2 target cells in lung cells of livestock, poultry, pets and wild animals

In consistent with previous report that SRAS-CoV-2 replicates poorly in chicken and duck^8^, no SARS-CoV-2 target cells were found in lung cells of poultry (chicken, duck, goose and pigeon) (Fig. 2a, b). In cat, SARS-CoV-2 target cells were detected in eight out of eleven cell types investigated, with the top two cell types being ciliated cells and secretory cells. In pig, co-expression of *ACE2* and *TMPRSS2* was observed in seven out of eleven cell types. In pangolin, a small proportion of endothelial cells were found to co-express *ACE2* and *TMPRSS2*. In hamster, ciliated cells were the cell type with most abundant SARS-CoV-2 target cells. In goat, we detected the co-expression of *ACE2* and *TMPRSS2* in ATI, fibroblasts, endothelial cells, ciliated cells and T cells. Goat share highly similar *ACE2* amino acids sequence with pig and human^22^, implying that goat *ACE2* might have similar capability for mediating virus entering into host cells. In lizard, we detected the co-expression of *ACE2* and *TMPRSS2*, mainly in B cells. In dog, less than 0.5% *ACE2* and *TMPRSS2* co-expressing cells were detected in ciliated cells and ATII (Fig. 2a, b).

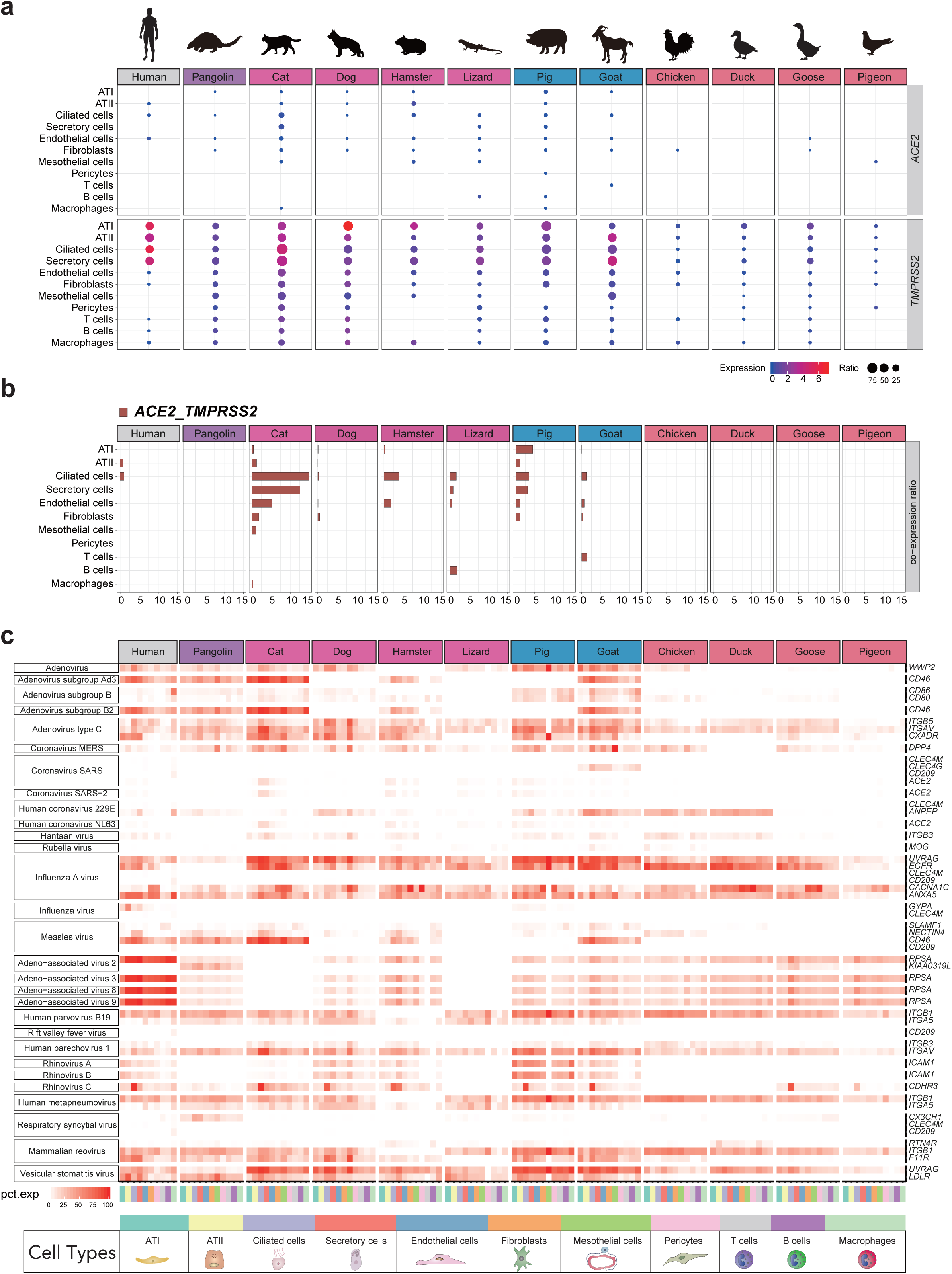

The infection with SARS-CoV-2 could lead to a severe pneumonia, and respiratory diseases caused by other respiratory viruses is also noteworthy. To reveal the putative target lung cells of other respiratory viruses, we screened the expression patterns of 32 virus receptors for a total of 29 virus species derived from 12 virus families (*coronavirida, orthomyxoviridae, adenoviridae, hantaviridae, matonaviridae, paramyxoviridae, parvoviridae, phenuiviridae, picornaviridae, pneumoviridae, reoviridae* and *rhabdoviridae*) (Fig. 2c, Supplementary Table 4), which have been shown to be able to transmit via the respiratory system^22^. Generally, poultry lung cells express less types of virus receptors than mammalians and reptiles. For example, coronavirus receptors were generally not expressed in poultry lung cells, except that human coronavirus 229E receptor *ANPEP* was expressed in chicken and duck lung cells but not in goose and pigeon lung cells. MERS coronavirus receptor *DPP4* was expressed in chicken and goose lung cells but was barely detected in corresponding cell types in duck and pigeon. Coronavirus SARS receptor *CLEC4G* was expressed in goat lung cells (ciliated cells, endothelial cells, fibroblasts and macrophages) but was found absent in lung cells of other species. Influenza A virus receptors *UVRAG* and *EGFR* was present in lung cells of every species we investigated, but was absent or exhibited meagre expression in pangolin and pigeon. *Adenoviridae* virus receptors showed a preferential expression in mammals, except adenovirus type C receptors which was also present in poultry lung cells. Rhinovirus C receptor displayed preferential expression in ciliated cells of human, cat, dog, hamster, pig, goat and goose while respiratory syncytial virus receptor *CD209* was only present in human macrophage cells. Taken together, our work, for the first time, revealed the putative target cells for respiratory viruses in an important organ of respiratory system (lung), which lays the foundation for dissecting the infection and transmission of respiratory system viruses at the single-cell level.

### Systematic evaluation of SARS-CoV-2 infection risks in livestock, poultry, pets and wildlife

While comparing the frequencies of SARS-CoV-2 target lung cells across different species, we noticed that cat clearly outweigh other species, with 13.19% in ciliated cells, compared to pig (3.35%) and hamster (3.87%) (Supplementary Table 3). Moreover, when taking the proportions of SARS-CoV-2 target cells in distinct organs among cat, pangolin and pig into consideration, it further indicated that the proportion of *ACE2* and *TMPRSS2* target cells were much higher in cat. For example, the proportion of *ACE2* and *TMPRSS2* co-expressing cell was as high as 40% in cat kidney proximal tubular cells while the proportions were only around 3% and 2% in corresponding cell type in pangolin and pig, respectively (Supplementary Table 3). We also noticed that SARS-CoV-2 target cells were widely distributed among organs within the digestive system (esophagus, rectum), respiratory system (lung) and urinatory system (kidney) of cat (Fig. 3a), implying that cats could be infected by SARS-CoV-2 via multiple routes such as dietary infections of the digestive tract or airborne transmission through respiratory system. To highlight SARS-CoV-2 susceptible cell types, we summarized all cell types with proportion of *ACE2* and *TMPRSS2* co-expressing cells higher than 1%, which clears shows that cat, as well as pig, have more susceptible cell types than other species we investigated (Fig. 3a). Taken together, our data explains the observation that cats are highly permissive to SARS-CoV-2^8^ and raise the necessity to carefully monitor and evaluate the possible roles of cat as intermediate hosts in current pandemic.

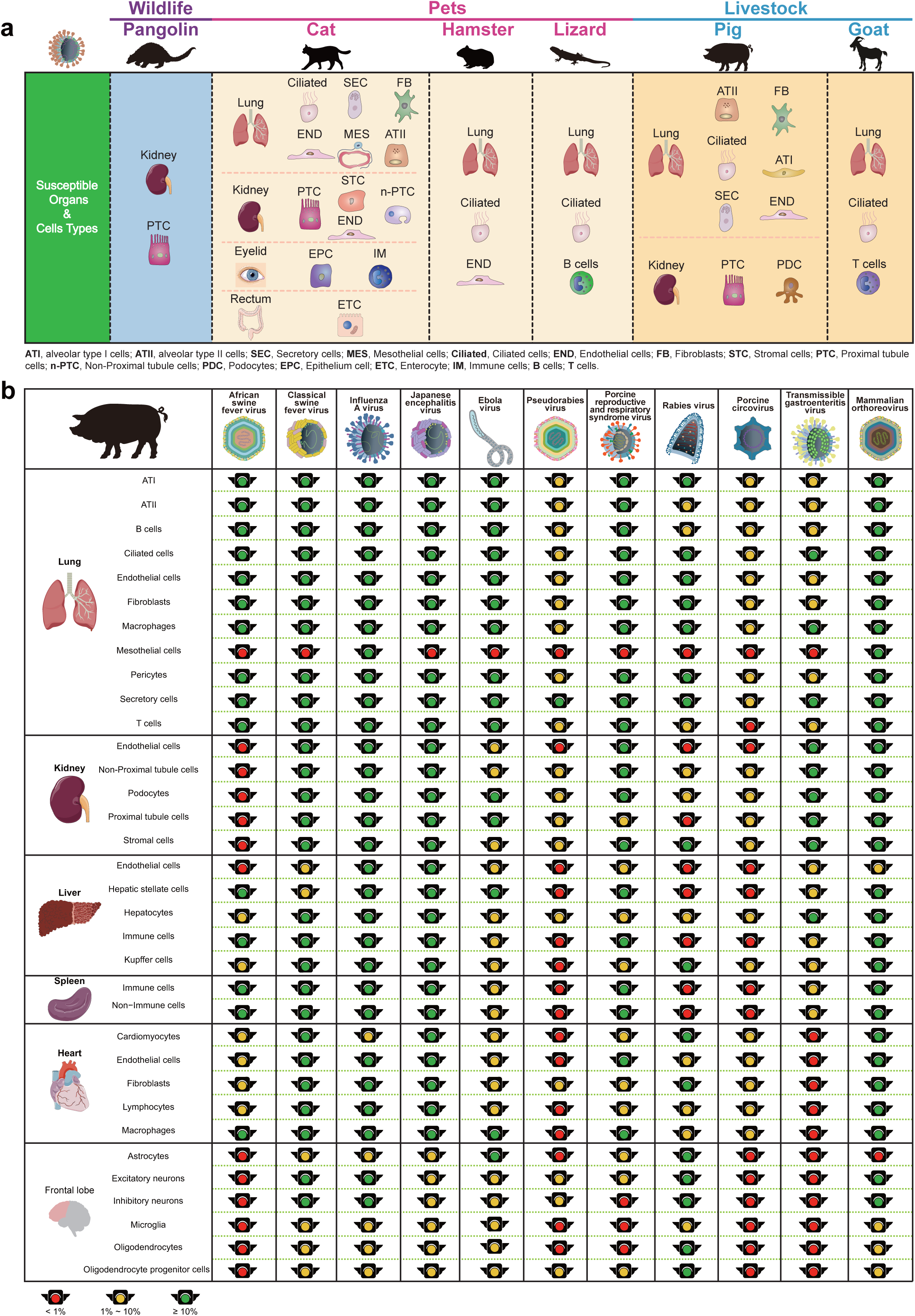

SARS-CoV-2 replicate poorly in dog, chicken and duck^8^. Our data suggests that the co-expression of *ACE2* and *TMPRSS2* is very rare in dog lung cells and absent in poultry lung cells (chicken, duck, goose and pigeon). Besides, it has been predicted that dog and poultry ACE2 cannot be utilized efficiently by SARS-CoV-2 spike glycoprotein because of mutations in critical amino acids of dog ACE2^5^. Therefore, our data, to some extent, explains why dogs and poultry are not as permissive for SARS-CoV-2 infection as cat.

In addition to cats, pangolins^6,7^ and hamsters^12^ have been reported to be permissive for SARS-CoV-2 infection, however, the target cells for virus infection and putative transmission routes are largely unknown. Our study identified the SARS-CoV-2 target cells in distinct tissues of cat, pangolin and hamster, indicated by the simultaneous expression of SARS-CoV-2 entry factors: *ACE2* and *TMPRSS2*. A detailed comparative analysis among cat, pangolin and hamster deciphered that proportion of SARS-CoV-2 target cells in cat was much higher than pangolin and hamster, implying that cats are more susceptible to SARS-CoV-2. Besides, as a companion animal, cats interact with humans more frequently than pangolins, thus we proposed that cats should be closely monitored in the current COVID pandemic. In addition, we also detected the co-expression of *ACE2* and *TMPRSS2* in lung cells of goat and lizard. Considering the lack of clear evidence showing weather they can be infected by SARS-CoV-2, it is noteworthy to evaluate their possibility to be the SARS-CoV-2 intermediate hosts.

Zhou *et al*. proved that pig ACE2 is capable of assisting SARS-CoV-2 entering into HeLa cells^23^, and we found that SARS-CoV-2 target cells were distributed in a variety of cells in the kidney and lung of domestic pig. Combining our analysis with the result from Zhou *et al*., it seems to be plausible to question if domestic pigs have the potential to function as intermediate hosts for SARS-CoV-2. However, in the experiments performed by Shi *et al*., pigs were found to be negative for viral RNA and antibody tests on day 2, day 4 and day 6 after inoculating intranasally with SARS-CoV-2^8^. The reason why pigs could not support SARS-CoV-2 replication^8^ seems to be an intriguing scientific question and definitely invites further investigations. Albeit, according to the current experimental evidence, pig does not seem to be permissive for SARS-CoV-2 infection and transmission, we need to be cautious about the putative roles of pig in future coronavirus outbreak, considering that pig expresses both SARS-CoV-2 entry factor (*ACE2*) and activator factor (*TMPRSS2*) in a variety of cell types and there was reported case of SARS-associated coronavirus transmission from human to pig^17^. Given that COVID-19 pandemic is still progressing and SARS-CoV-2 strains are constantly evolving^24^, we need to keep monitoring and evaluating the possibility of pigs to become intermediate hosts of future pandemic.

### Systematically screening of target cells for 144 viruses

To investigate the susceptibility of host cells to different kinds of viruses, we screened the expression patterns of 114 receptors of 144 viruses (representing 26 virus families) in distinct organs of cat, pangolin and pig and the lung of pets, livestock, poultry and wildlife in an unbiased manner, resulting in a comprehensive atlas of virus target cells (Extended Data Fig. 5-7, Supplementary Data Table 5). Intriguingly, we found Rabies lyssavirus receptor *NCAM1* was widely expressed in cell types of pig neural system but rarely expressed in non-neural tissues. Besides, we found that vesicular stomatitis virus receptor *UVRAG* was preferentially expressed in pig lung cells (Extended Data Fig. 6). To demonstrate the proportions of virus receptor expressing cells in an intuitive manner, we proposed a system named “traffic light system for virus entry” to assign each cell type to one of the following three status: red light, green light and yellow light. Briefly, if the receptor is expressed in less than 1% of that cell types, a “red light” status will be assigned. If the proportion is between 1% and 10%, “yellow light” status will be assigned. If the proportion is over 10%, then a “green light” status would be assigned (see methods). Based on this system, we assigned status to all the cell populations of all the investigated species (Supplementary Data Table 5). To demonstrate the potential usefulness of this system, we employed this analysis and visualized the status of several representative viruses in distinct organs of pig (Fig. 3b and Extended Data Fig. 7), cat (Supplementary Data Fig. 5) and pangolin (Extended Data Fig. 6) respectively. Overall, our project provides a model framework for future research about the screening of host susceptibility, and effectively demonstrates the applicability of single cell atlas resources in exploring the expression patterns of virus receptors and led to the identification of putative virus target cells.

Host susceptibility has been evaluated using *in vitro* assay or *in vivo* inoculation experiments under laboratory circumstance, which cannot fully recapitulate or simulate the real process where virus co-evolve with hosts. Besides, it only reflects the susceptibility of a host to a specific virus under current situation and fail to consider the dynamic interaction process between viruses and hosts. Considering the substantial mutation rates and adaptation abilities of viruses^25^, it is important to evaluate virus infection and transmission capability in a more fundamental manner. Viruses and hosts have been constantly co-evolving for millions of years, which collectively shaped the immune landscape of animals and the host range of viruses^25^. Virus receptors are the key determining factors for virus entry, thus dissecting virus receptor expression pattern is fundamental for understanding the intra-species and inter-species transmission of viruses, both in current and future infectious diseases.

Our project provides a novel strategy to find out putative susceptible hosts, based on the distribution of virus target cells among distinct cell populations, which makes it possible to screen the susceptibility of all existing viruses (using the available virus receptor information) on all existing species (using the available scRNAseq data) in an unbiased manner. With the development of single cell sequencing techniques and the progress of international single cell atlas collaborative projects, the atlas for more species could be generated at an accelerated speed. We anticipate that the information gained from the present study will certainly augment future research work, and provide some novel insights about the prevention and control strategies against SARS-CoV-2 along with many other harmful viruses.

## Materials and methods

### Ethics statement

Sample collection and research were performed with the approval of Institutional Review Board on Ethics Committee of BGI (Approval letter reference number BGI-NO. BGI-IRB A20008). All procedures were conducted according to the guidelines of Institutional Review Board on Ethics Committee of BGI.

### Sample collection

A total of 11 samples were collected in this study, including four pets: *Felis catus* (cat), *Canis lupus familiaris* (dog), *Mesocricetus auratus* (hamster), *Anolis carolinensis* (lizard), two livestock: *Sus scrofa domesticus* (pig), *Capra aegagrus hircus* (goat), four poultry: *Gallus gallus domesticus* (chicken), *Anser cygnoides domesticus* (goose), *Anas platyrhynchos domesticus* (duck), *Columba livia domestica* (pigeon), and one wild animal: *Manis javanica* (pangolin). The pets were bought from a pet market, and the livestock and poultry were purchased from an agricultural market. The *Manis javanica* sample was collected from a pangolin which died of natural causes in Guangdong Provincial Wildlife Rescue Center and immediately stored in −80°C freezer after dissection.

After the execution of animals in accordance with the ethics of animal experiment, dissection was carried out quickly to separate each organ. The collected tissues were rinsed by 1X PBS, then quick-frozen and stored in liquid nitrogen. The single cell nucleus of each tissue was separated by mechanical extraction method^26^. Briefly, the tissues were first thawed, infiltrated by 1X homogenization buffer (containing 30mM CaCl_2_, 18mM Mg(Ac)_2_, 60mM Tris-HCl (pH 7.8), 320mM sucrose, 0.1% NP-40, 0.1mM EDTA and 0.2U/µl RNase inhibitor), then cut into smaller pieces, and the single nucleus was isolated by 2ml Dounce homogenizer set. After filtration with 30µm strainer, the nuclei extraction was resuspended by 1%BSA containing 0.2U/µl RNase inhibitor and span down at the speed of 500g for 10 min at 4 degrees (to carefully discard the cellular impurities within the supernatant). This step was repeated twice, and finally the nucleus was recollected with 0.1% BSA containing 0.2U/µl RNase inhibitor. Subsequently, DAPI was used to stain the nucleus, and the nucleus density was calculated under a fluorescence microscope to prepare for the subsequent library construction.

### Single nuclei library construction and sequencing

The mRNA within the single nucleus samples of different organs of pig (heart, liver, spleen, lung, kidney, hypothalamus, area postrema, vascular organ of lamina terminalis, subfomical organ and cerebellum) were captured and the libraries were constructed using inhouse DNBelab C4 kit and sequenced using DNBSEQ-T1. The separated single nucleus of different organs (including the lungs for pig, dog, cat, goat, pangolin, chicken, pigeon, goose, duck, lizard, and hamster; pangolin organs: heart, liver, spleen, lung, kidney, large intestine, duodenum, stomach and esophagus; cat organs: heart, liver, lung, kidney, eyelid, esophagus, duodenum, colon and rectum) were constructed using Chromium Single Cell 3LJ GEM, Library & Gel Bead Kit v3 (PN-1000075) following the standard user guide provided by manufacturer. After performing the library conversions using the MGIEasy Universal DNA Library Preparation Reagent Kit, the libraries were sequenced by compatible BGISEQ-500 sequencing platform.

### Cross-species homolog conversion

To facilitate the integration of cross-species lung single cell data set, we converted genes from other species to mouse homologs. We downloaded homolog gene lists using BioMart^27^. Then, if a 1:1 match existed between a non-mouse gene and a mouse gene, the non-mouse gene name was converted. As for pangolin, goose and pigeon, which lack homologs records on Ensemble, single-copy orthologs were identified from two species genomes by cluster analysis of gene families using OrthoFinder^28^ (OrthoFinder version 2.3.3) with the default parameters.

### Single-cell RNAseq data processing

Sequencing data filtered using custom script and gene expression matrix were obtained using Cell Ranger 3.0.2 (10X Genomics). The genomes using for reads alignment were downloaded from NCBI Assembly (Supplementary Table 6). Single cell analysis was conducted using Seurat^19,29^. Briefly, quality control was performed based on the following criteria: cells with mapped number of genes less than 200 or with mitochondrial percentage higher than 10% were removed. Variable genes were determined using Seurat’s FindVariableGenes function with default parameters. Clusters were identified using Seurat’s FindClusters function and visualized using Seurat’s RunTSNE. All the DEGs for each Seurat Objects were identified using Seurat’s FindAllMarkers function. Cell types were annotated according to the expression of canonical cell type markers.

### Integration of lung data sets from different species

The human lung single cell RNAseq data was obtained from literature^30^. Data sets of lungs from different species were integrated using Seurat’s FindIntegrationAnchors and IntegrateData function with features after homolog conversion.

### Virus receptor list collection

Virus receptor list were downloaded from a virus–host receptor interaction database^31^ and manually collected from published literatures (Supplementary Table 7).

### Domestic pig brain data set collection

Single nuclei RNAseq data sets for frontal lobe, occipital lobe, parietal lobe, temporal lobe and hypothalamus of domestic pig were obtained from literature^26^.

### Traffic light status assignment

The proportions of virus receptor were calculated and “red, yellow, green” status was assigned to each cell type based on receptor proportions (less than 1%, red light; greater than or equals to 1% & less than 10%, yellow light; greater than or equals to 10%, green light). In case of viruses with multiple receptors, the receptor with the highest proportion was considered for status assignment.

## Data availability

The single cell atlas of all the investigated species in this study are available via http://120.79.46.200:81/SARS-CoV-2. The raw data supporting the findings of this study will be made available upon request. Raw transcriptome sequencing data has been deposited to the CNSA (CNGB Nucleotide Sequence Archive) with the accession number CNP0001085 (https://db.cngb.org/cnsa/) and will be released to the public after the manuscript is accepted for publication.

## Acknowledgement

This work was supported by China National GeneBank (CNGB). Our project was financially supported by funding from the National Key R&D Program of China (No. 2019YFC1711000), National Key Research and Development Program of China (2016YFD0501300), the Program for Innovative Research Team in the University of Ministry of Education of China (IRT_17R39), the Guangdong Provincial Key Laboratory of Genome Read and Write (grant No. 2017B030301011). We thank the Guangdong Provincial Academician Workstation of BGI Synthetic Genomics, BGI-Shenzhen, Guangdong, China. We thank Professor Wensheng Zhang (Suzhou University) and Professor Gang Cao (Huazhong Agricultural University) for creative discussion and informative feedback. Finally, we are thankful to the China National GeneBank for producing the sequencing data.

## Author contributions

XX, HL, YHL, DSC, JS, YH, HMY, WQ, LNZ conceived and designed the project. JCZ, WYW, HRL, FYA, HB, ZHC, QH were responsible for sample collection and dissection. JCZ, WYW, HRL participated in single nuclei library construction and sequencing. DSC, JCZ, XND, TML coordinated data analysis. RX, XLW, HYW, PWD, JYQ, SYW, HML, LZ, LH, YXZ, HXC, DMF, LHL, YYZ, FYW, YTY, WDW, CCS, XRW contributed to single cell clustering and cell annotation. XX, HL, YHL, DSC, JS, YH, HMY, WQ, YQW, JHW, XHS, SJJ, LC, SKS, SHZ, XYM, WMF, YX, FL, ZML, FC, XJ participated in data interpretation, visualization and manuscript writing.

---

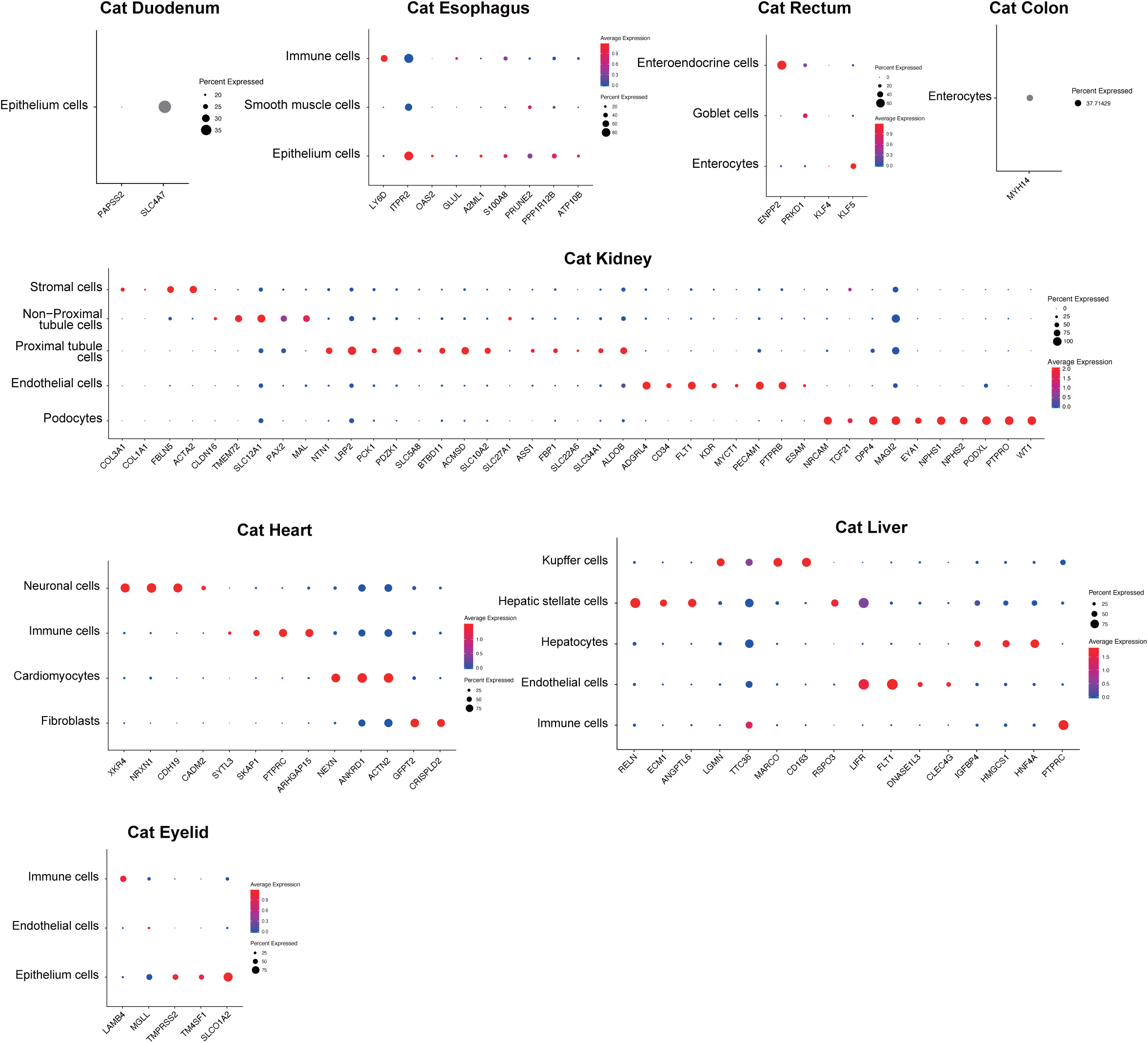

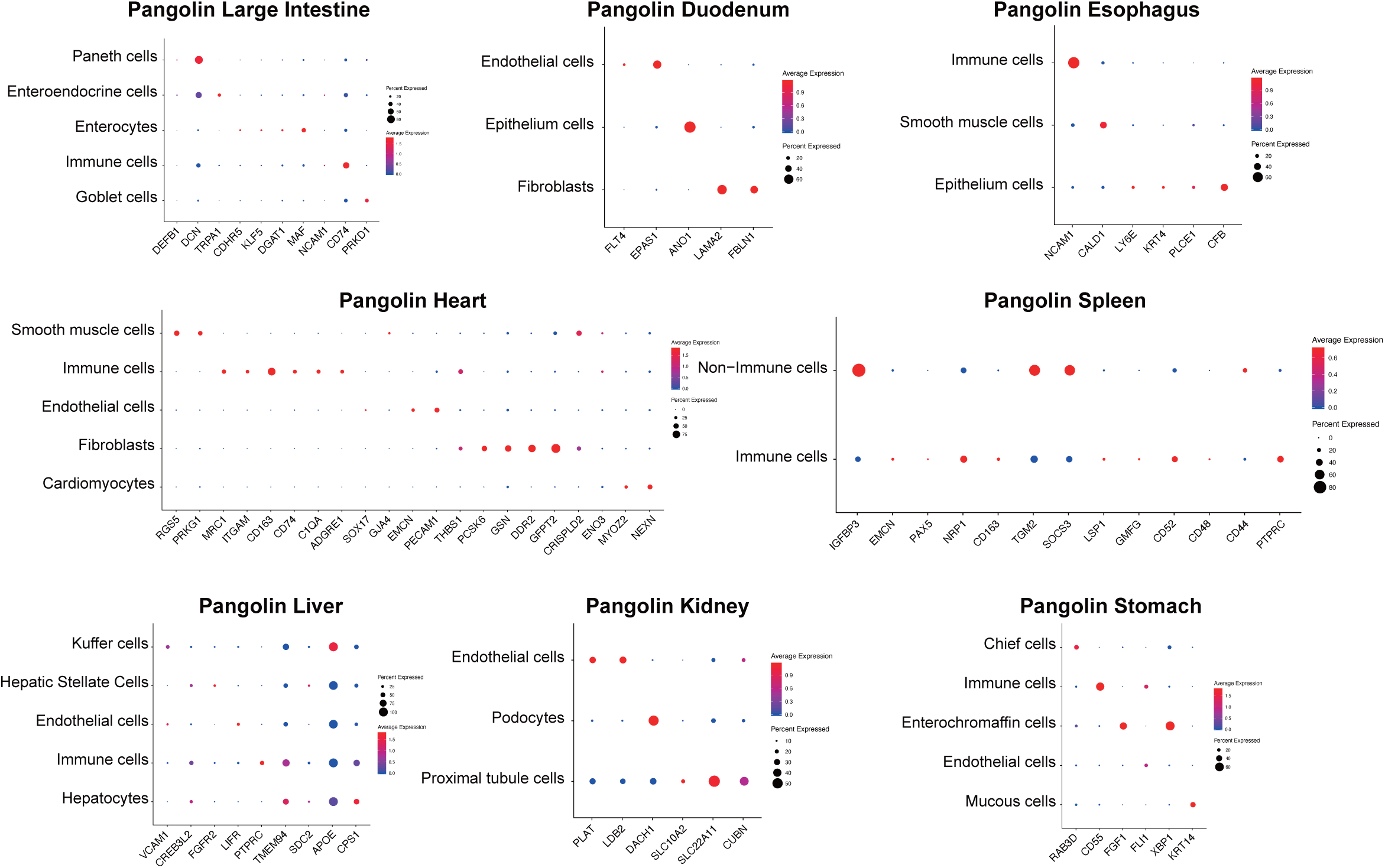

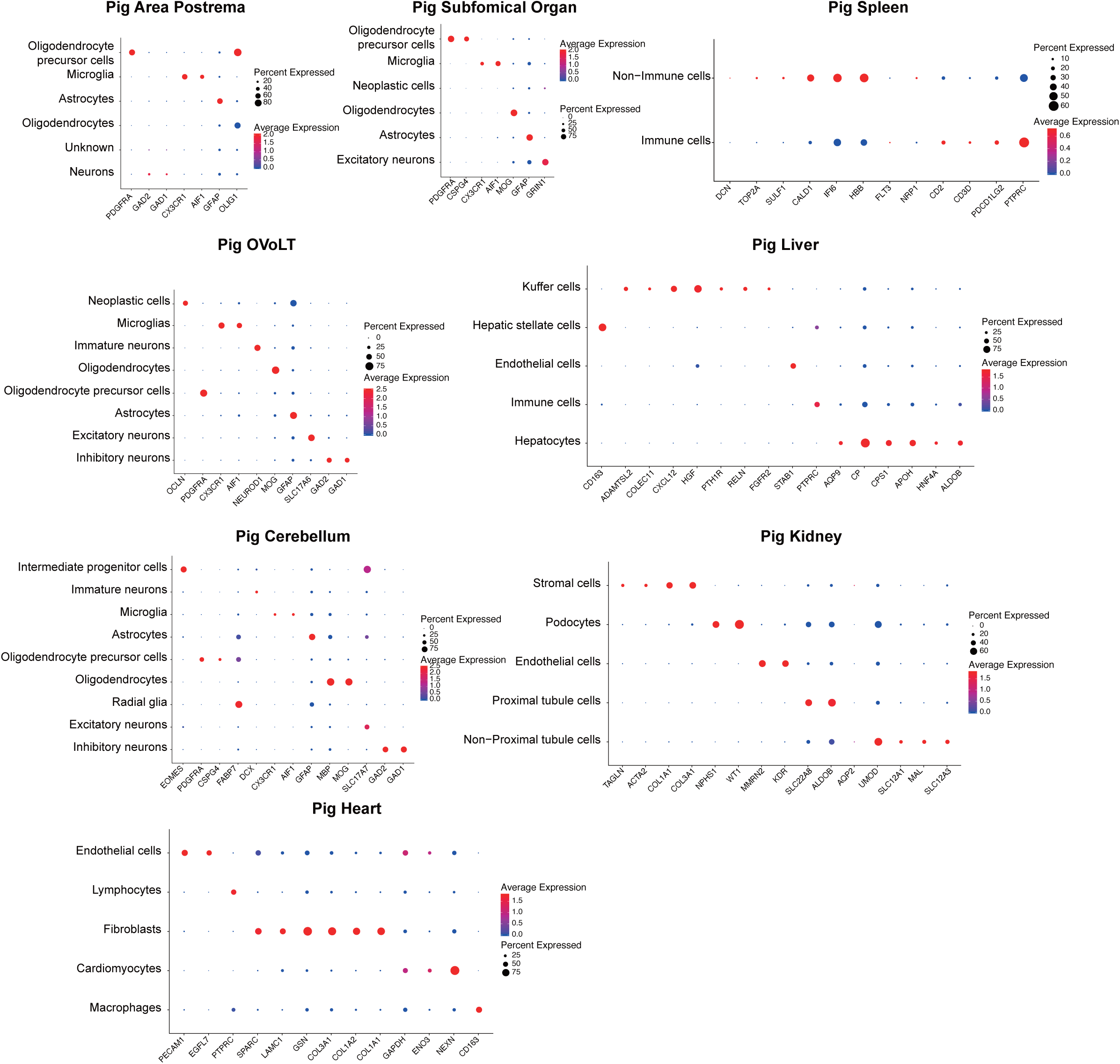

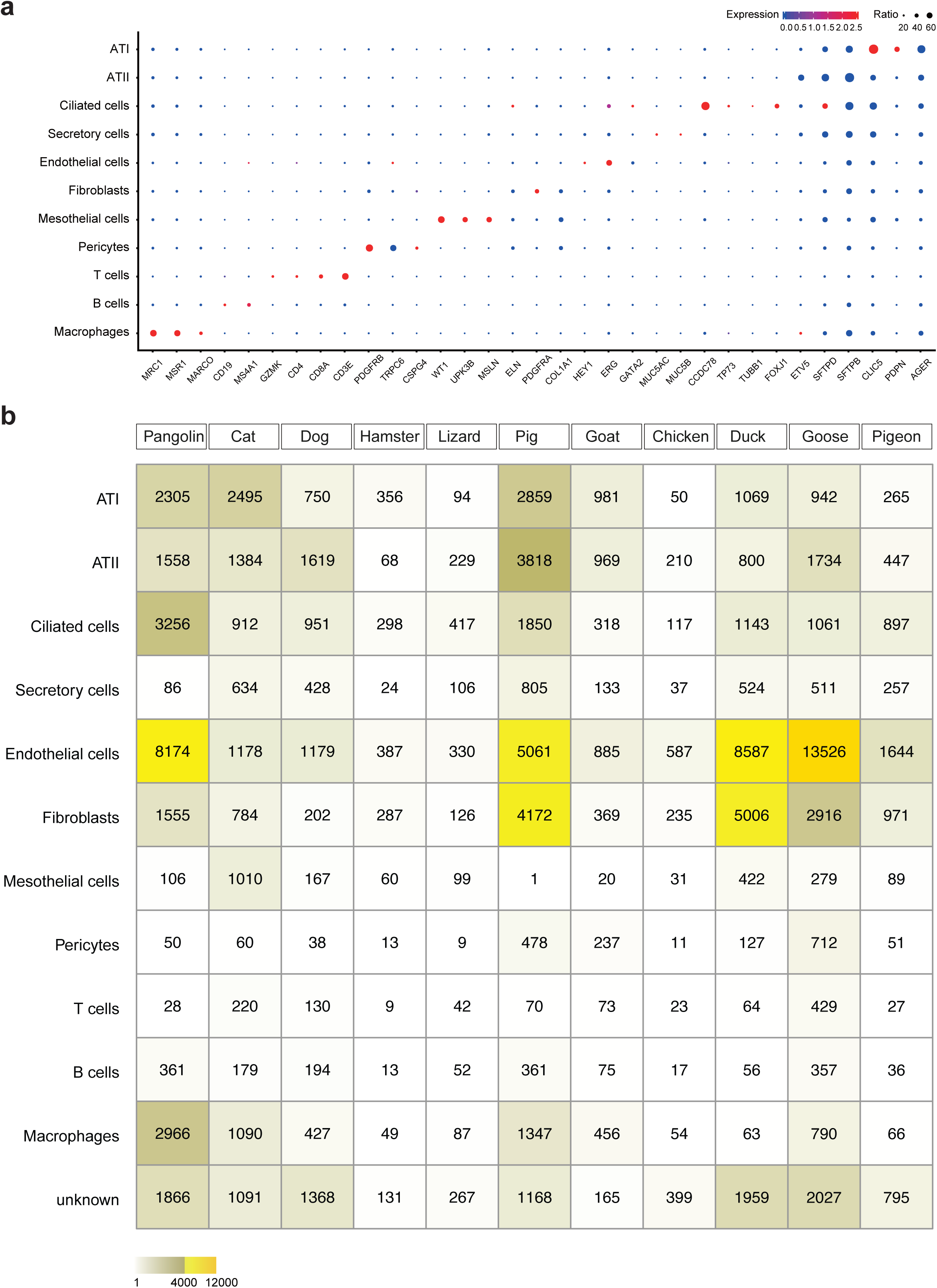

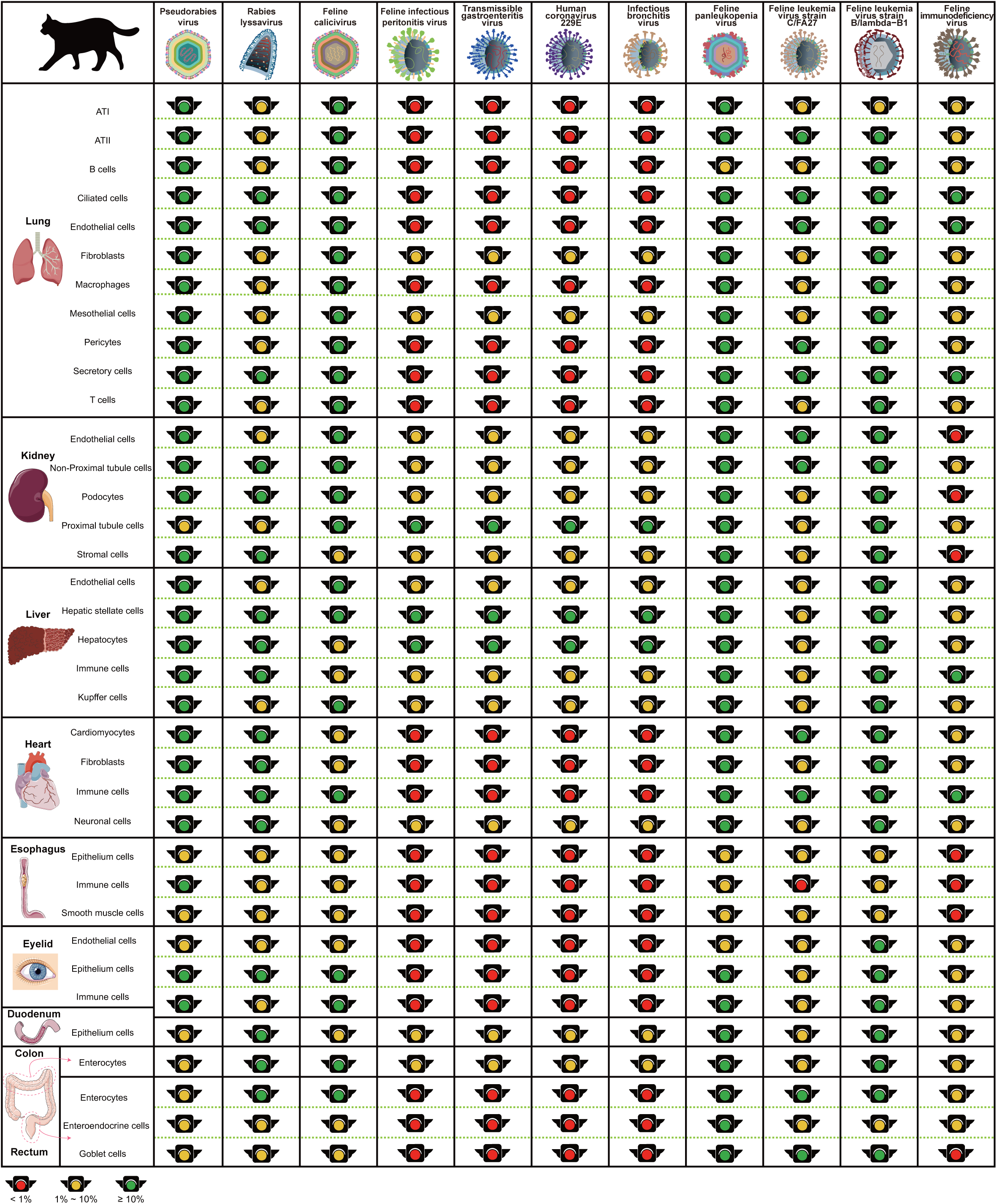

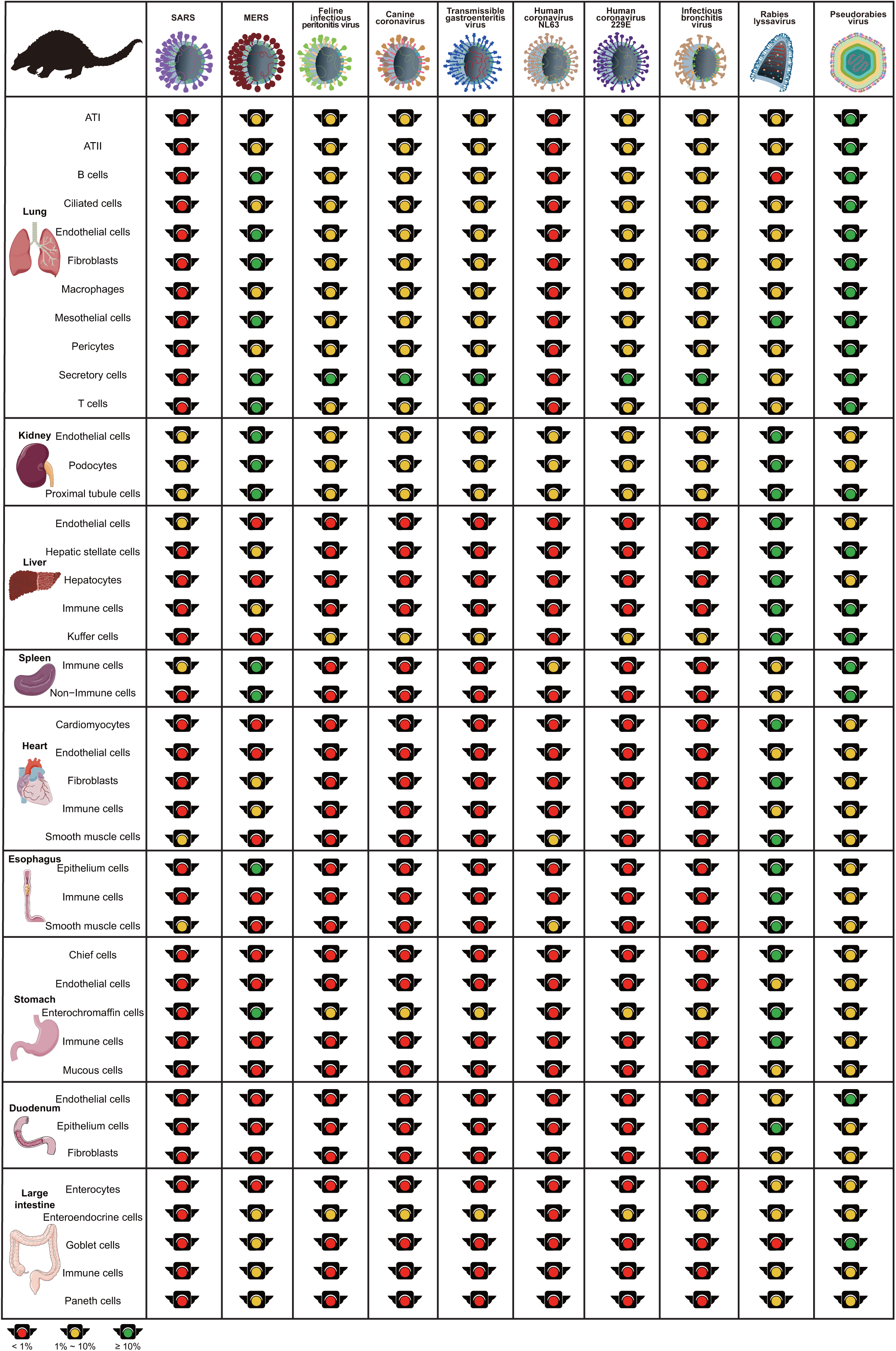

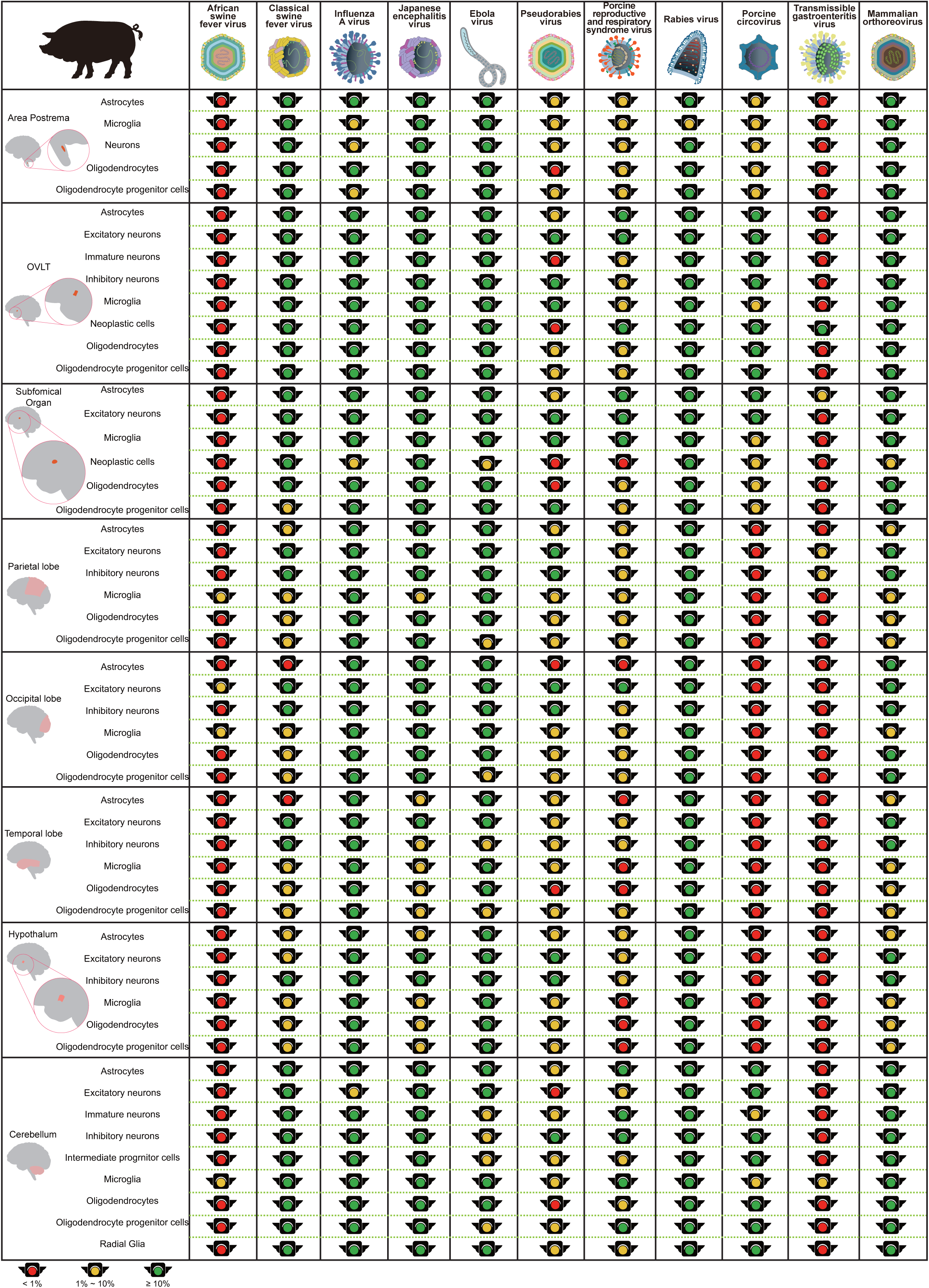

## References

1. De Wit, E., Van Doremalen, N., Falzarano, D. & Munster, V. J. SARS and MERS: Recent insights into emerging coronaviruses. Nature Reviews Microbiology vol. 14 523–534 (2016).

2. Baud, D., Gubler, D. J., Schaub, B., Lanteri, M. C. & Musso, D. An update on Zika virus infection. The Lancet vol. 390 2099–2109 (2017).

3. Smith, G. J. D. et al. Origins and evolutionary genomics of the 2009 swine-origin H1N1 influenza a epidemic. Nature 459, 1122–1125 (2009).

4. COVID-19 situation reports. https://www.who.int/emergencies/diseases/novel-coronavirus-2019/situation-reports/.

5. Lau, S. K. P. et al. Possible Bat Origin of Severe Acute Respiratory Syndrome Coronavirus 2. Emerg. Infect. Dis. 26, (2020).

6. Lam, T. T. Y. et al. Identifying SARS-CoV-2 related coronaviruses in Malayan pangolins. Nature 1–6 (2020) doi:10.1038/s41586-020-2169-0.

7. Xiao, K. et al. Isolation of SARS-CoV-2-related coronavirus from Malayan pangolins. Nature 1–7 (2020) doi:10.1038/s41586-020-2313-x.

8. Shi, J. et al. Susceptibility of ferrets, cats, dogs, and other domesticated animals to SARS–coronavirus 2. Science (80-.). eabb7015 (2020) doi:10.1126/science.abb7015.

9. Zhang, Q. et al. SARS-CoV-2 neutralizing serum antibodies in cats: a serological investigation. bioRxiv 2020.04.01.021196 (2020) doi:10.1101/2020.04.01.021196.

10. Halfmann, P. J. et al. Transmission of SARS-CoV-2 in Domestic Cats. N. Engl. J. Med. NEJMc2013400 (2020) doi:10.1056/NEJMc2013400.

11. Sit, T. H. C. et al. Infection of dogs with SARS-CoV-2. Nature 1–6 (2020) doi:10.1038/s41586-020-2334-5.

12. Sia, S. F. et al. Pathogenesis and transmission of SARS-CoV-2 in golden hamsters. Nature 1–7 (2020) doi:10.1038/s41586-020-2342-5.

13. Hoffmann, M. et al. SARS-CoV-2 Cell Entry Depends on ACE2 and TMPRSS2 and Is Blocked by a Clinically Proven Protease Inhibitor. Cell 181, 271-280.e8 (2020).

14. Matsuyama, S. et al. Efficient Activation of the Severe Acute Respiratory Syndrome Coronavirus Spike Protein by the Transmembrane Protease TMPRSS2. J. Virol. 84, 12658–12664 (2010).

15. Regev, A. et al. The human cell atlas. Elife 6, (2017).

16. Sungnak, W. et al. SARS-CoV-2 entry factors are highly expressed in nasal epithelial cells together with innate immune genes. Nat. Med. 26, 681–687 (2020).

17. Chen, W. et al. SARS-associated coronavirus transmitted from human to pig. Emerg. Infect. Dis. 11, 446–448 (2005).

18. Butler, A., Hoffman, P., Smibert, P., Papalexi, E. & Satija, R. Integrating single-cell transcriptomic data across different conditions, technologies, and species. Nat. Biotechnol. 36, 411–420 (2018).

19. Stuart, T. et al. Comprehensive Integration of Single-Cell Data. Cell 177, 1888-1902.e21 (2019).

20. Ziegler, C. et al. SARS-CoV-2 Receptor ACE2 is an Interferon-Stimulated Gene in Human Airway Epithelial Cells and Is Enriched in Specific Cell Subsets Across Tissues. SSRN Electronic Journal http://shaleklab.com/publication/sars-cov-2-receptor-ace2-is-an-interferon-stimulated-gene-in-human-airway-epithelial-cells-and-is-detected-in-specific-cell-subsets-across-tissues/ (2020) doi:10.2139/ssrn.3555145.

21. Tay, M. Z., Poh, C. M., Rénia, L., MacAry, P. A. & Ng, L. F. P. The trinity of COVID-19: immunity, inflammation and intervention. Nat. Rev. Immunol. 1–12 (2020) doi:10.1038/s41577-020-0311-8.

22. Qiu, Y. et al. Predicting the angiotensin converting enzyme 2 (ACE2) utilizing capability as the receptor of SARS-CoV-2. Microbes Infect. (2020) doi:10.1016/j.micinf.2020.03.003.

23. Zhou, P. et al. A pneumonia outbreak associated with a new coronavirus of probable bat origin. Nature 579, 270–273 (2020).

24. Forster, P., Forster, L., Renfrew, C. & Forster, M. Phylogenetic network analysis of SARS-CoV-2 genomes. Proc. Natl. Acad. Sci. 117, 202004999 (2020).

25. Duxbury, E. M. L. et al. Host-pathogen coevolution increases genetic variation in susceptibility to infection. Elife 8, (2019).

26. Chen, D. et al. Single cell atlas of domestic pig brain illuminates the conservation and divergence of cell types at spatial and species levels. bioRxiv 2019.12.11.872721 (2019) doi:10.1101/2019.12.11.872721.

27. Cunningham, F. et al. Ensembl 2019. Nucleic Acids Research vol. 47 D745–D751 https://www.ncbi.nlm.nih.gov/pmc/articles/PMC6323964/ (2019).

28. Emms, D. M. & Kelly, S. OrthoFinder: Phylogenetic orthology inference for comparative genomics. Genome Biol. 20, 238 (2019).

29. Butler, A., Hoffman, P., Smibert, P., Papalexi, E. & Satija, R. Integrating single-cell transcriptomic data across different conditions, technologies, and species. Nat. Biotechnol. 36, 411–420 (2018).

30. Raredon, M. S. B. et al. Single-cell connectomic analysis of adult mammalian lungs. Sci. Adv. 5, eaaw3851 (2019).

31. Zhang, Z. et al. Cell membrane proteins with high N-glycosylation, high expression and multiple interaction partners are preferred by mammalian viruses as receptors. Bioinformatics vol. 35 723–728 https://pubmed.ncbi.nlm.nih.gov/30102334/ (2019).

