## Supplementary material for "Single-cell screening of SARS-CoV-2 target cells in pets, livestock, poultry and wildlife": Legend.

**Supplementary Table 1|** **Quality control information of single nuclei sequencing libraries constructed using Chromium Single Cell 3ʹ GEM, Library & Gel Bead Kit (10X) and inhouse DNBelab C4 kit respectively.**

**Supplementary Table 2|** **Differentially expressed genes (DEGs) of each cluster within each tissue of pig, pangolin, cat and lungs of various species.**

**Supplementary Table 3|** **Detailed expression percentage of *ACE2*, *TMPRSS2* and co-expression of two genes within each cell type of various species.**

**Supplementary Table 4| Detailed expression percentage of virus receptors for respiratory viruses.**

**Supplementary Table 5| Detailed expression percentage of the total of 114 receptors from 144 viruses in each cell types of different species.**

**Supplementary Table 6| Reference genomes assemblies information. The genomes were downloaded from NCBI Assembly and used for reads alignment.**

**Supplementary Table 7| Virus receptor list collected from a virus-host receptor interaction database and published literatures.**

**Extended Data Fig. 1|** **Dot plots showing expression patterns of cell type marker genes in different tissues of cat.**

**Extended Data Fig. 2|** **Dot plots showing expression patterns of cell type marker genes in different tissues of pangolin.**

**Extended Data Fig. 3|** **Dot plots showing expression patterns of cell type marker genes in different tissues of pig.**

**Extended Data Fig.4| Cell type annotation of comparative lung atlas. a**, Dot plots showing expression patterns of cell type marker genes in lung cells of different species. **b**, The distribution of lung cells within each species.

**Extended Data Fig.5|Traffic light system showing virus infection risks in different cell types across tissues in cat.**

**Extended Data Fig.6|Traffic light system showing virus infection risks in different cell types across tissues in pangolin.**

**Extended Data Fig.7|Traffic light system showing virus infection risks in different cell types across tissues in pig.**
